# Children Develop Adult-Like Visual Sensitivity to Image Memorability by the Age of Four

**DOI:** 10.1101/2022.12.20.520853

**Authors:** Xiaohan (Hannah) Guo, Wilma A. Bainbridge

## Abstract

Adults have been shown to consistently remember and forget certain images despite large individual differences, suggesting a population-wide sensitivity to an image’s intrinsic *memorability*—a measure of how successfully an image is remembered. While a decade of research has focused on image memorability among adults, the developmental trajectory of these consistencies in memory is understudied. Here, we investigate by what age children gain adultlike sensitivity to the image memorability effect. We utilized data from Saragosa-Harris et al. (2021), where 137 children aged between 3 and 5 years old encoded animal-scene image pairs and then after a 5-minute, 24-hour, or 1-week delay performed a cued recognition task for each scene target given its animal cue. We tested adults’ memory of the same scene images using ResMem (Needell & Bainbridge, 2022), a pre-trained deep neural network that predicts adult image memorability scores, and using an online behavioral continuous recognition task *(N* = 116). Results showed that ResMem predictions, as a proxy of adults’ memory, predicted scene memory of children by the age of 4 and were the most predictive of children’s memory across ages after a long, 1-week delay. Children at age 3 show non-adult-like consistent memory patterns, implying that the non-adult-like memory patterns were not due to poor memory performance. Instead, 3-year-olds may have consistently used certain visual memory strategies that become less optimal as they age. Our results suggest that adult-like sensitivity to image memorability emerges by the age of 4 through experience.

**Public Significance Statement:** This study strongly suggests that children older than 4 years old tend to remember and forget the same images as adults. We recommend teachers and caregivers to utilize the ResMem DNN to select memorable images to be used in educational settings.

## Introduction

We experience a continuous, endless stream of visual sensory information daily, where some visual information is remembered, and other information is forgotten. Despite large individual differences in personal experience and memory performance, there is an overwhelming consistency in what visual information adults tend to remember and forget (Bainbridge et al., 2013; Isola et al., 2011). Indeed, some scenes (Isola et al., 2011), faces (Bainbridge et al., 2013), words (Xie et al., 2020), videos (Cohendet et al., 2018), and even dance movements (Ongchoco et al., 2022) are intrinsically more memorable than others. This phenomenon whereby adults consistently remember and forget certain visual stimuli can be captured by *memorability* (Bainbridge et al., 2013; Isola et al., 2011), an objective stimulus measure indicating how well observers remember certain sensory information, across the adult population. How and when the susceptibility to memorability emerges across development, however, is understudied. It remains unclear whether children also consistently remember and forget certain visual stimuli similar to adults, and if so, whether memorability scores derived from adults can be used to predict children’s memory patterns. In this study, we ask how and when in childhood humans develop adult-like consistent visual memory patterns.

One hypothesis is that humans develop susceptibilities to memorability in early childhood, driven by the development of specific brain structures or the accumulation of visual experience. Human and animal research together has highlighted a critical period in early childhood when the hippocampus undergoes a decline in neurogenesis and an increase in competence to store long-term memories (Akers et al., 2014; Alberini & Travaglia, 2017; Dennis et al., 2016; Josselyn & Frankland, 2012; Travaglia et al., 2016). Such neural development eliminates infantile amnesia, the phenomenon whereby episodic memories in early childhood quickly fade away and are rarely remembered in adulthood (Hayne, 2004). In humans, infantile amnesia fades away at 3 to 4 years of age (Bauer et al., 2011; Schneider & Pressley, 2013), implying a change in children’s memory from non-adult-like to more adult-like in this period due to rapid hippocampal development. Human functional magnetic resonance imaging (fMRI) research has found that the medial temporal lobe including the hippocampus may underlie the memorability effect (Bainbridge et al., 2017; Bainbridge & Rissman, 2018). Thus, it is possible that children develop a susceptibility to the memorability effect along with the resilience to infantile amnesia in the period of hippocampal development at ages 3 and 4. In addition, the accumulation of visual experience in the first few years of life could gradually train children’s visual memory to become more adult-like. Memorability may reflect the statistical properties of stimuli in our world (Hovhannisyan et al., 2021), such as co-occurrence statistics (Xie et al., 2020) or across-item similarity (Lukavský & Děchtěrenko, 2017, although see Kramer et al., 2022). Thus, consistencies in memory could develop as children learn these statistical regularities of visual stimuli.

An alternative hypothesis is that children possess adult-like susceptibility to the memorability effect early on, and we would expect young children to have similar memory patterns to adults. Previous research has found that infants are born with capabilities to process faces (Farah et al., 2000; Tzourio-Mazoyer et al., 2002), scenes (Kamps et al., 2020), and edges (Kessen et al., 1972). These commonalities in visual processing between newborns and adults indicate early sensitivity to certain visual categories and properties, occurring in late visual areas in the brain (i.e., the inferotemporal cortex). Magnetoencephalography (MEG) studies have found that adults’ sensitivity to memorability occurs rapidly at the time-scale of late visual processing (170-250 ms; Khaligh-Razavi et al., 2016; Mohsenzadeh et al., 2019), and an fMRI study found adult sensitivity to memorability in inferotemporal cortex (Bainbridge et al., 2017). Thus, like their early sensitivity to visual categories, it is possible that infants could possess sensitivity to memorability in late visual areas from birth. Another piece of supportive evidence for early sensitivity to the visual memorability effect lies in primate research. Despite deprived visual experience compared to humans, laboratory-reared monkeys showed higher magnitudes of brain activity in late visual areas when viewing real-world images (e.g., fire hydrant, grocery store) that were more memorable to humans (Jaegle et al., 2019). Such commonalities in visual processing across species suggest that limited visual experience is required to develop a susceptibility to the memorability effect, providing supportive evidence that infants could have similar visual sensitivity to adults and thus exhibit adult-like visual memory patterns very early on.

One power of memorability is that memorability score is highly consistent across different tasks and observers (Bainbridge, 2020; Goetschalckx et al., 2019). The classic method of computing memorability scores is to use a continuous recognition task, where a timed sequence of stimuli is shown, and participants are asked to indicate stimulus repeats (Bainbridge et al., 2013; Isola et al., 2011). Another way of computing memorability scores is to use a paired association task, where a timed sequence of stimulus pairs is encoded, and after a delay, participants are asked to indicate the target stimulus based on its paired counterpart as a cue (Xie et al., 2020). Previous results suggest that the memorability of the target stimulus rather than the cue drives memory performance. Memorability can also be measured from one adult population to successfully predict the memory of another adult population (e.g., from online participants to patients with intracranial electrodes; Xie et al., 2020). Thus, memorability can be measured in one group or task and used to predict memory in another group or task, and we can apply this principle to predicting children’s memory from adult behavior. It is also possible to predict memorability scores using a pre-trained deep neural network called ResMem (Needell & Bainbridge, 2022), which can significantly predict image memorability for adults (a correlation of 0.67 between predicted and true memorability scores). Knowing that memorability scores obtained across paradigms robustly reflect adults’ memory, in the current study, we intend to test children’s development of adult-like visual memory using itemized memorability obtained using neural network predictions and adult memory behavior.

The current study serves as the first study to investigate the development of sensitivity to image memorability, investigating the two previously mentioned hypotheses. We ask by what age children have developed adult-like visual sensitivity to the image memorability effect. We ran a series of experiments comparing children’s memory with adult memory measures for a set of scene images, and found that children develop adult-like sensitivity to image memorability by the age of 4. This memory pattern was best predicted by ResMem predictions of adult memorability, when the children were tested after a 1-week delay. Further, we found evidence that children at the age of 3 consistently used certain encoding strategies that were not applied by the older age groups. Previous research has shown that young children have less accurate episodic memory (Zimmermann & Meier, 2006) and worse recognition memory performance (Lorsbach & Reimer, 2005) compared to adults. Despite children’s worse memory, our results imply that at least one aspect of memory—the sensitivity to the memorability of images—becomes adult-like in early childhood while memory is still developing. Further, we have identified a deep neural network able to predict children’s memory, empowering educators and caregivers with a tool able to create more intuitive learning materials for young children.

## Method

### Child Memory Dataset

We utilized children’s image memory data from Saragosa-Harris et al. (2021) (OSF link: https://osf.io/4m9kx/), where 3- to 5-year-old children encoded and retrieved animal-scene image pairs in an associative memory task. We measured children’s memory accuracy (*percent correct*) for each scene and animal image using these memory test data.

In the original experiment (Saragosa-Harris et al., 2021), a sample of 137 children (*N* female = 72) aged 3 to 5 years old (45 3-year-olds, 45 4-year-olds, and 47 5-year-olds) first encoded eight animal-scene associative pairs and was then tested with a cued recognition task after a delay. Child participants in each age group were evenly assigned to three delay conditions, where memory was assessed after a 5-minute (*N* = 45; 15 3-year-olds, 15 4-year-olds, and 15 5-years-olds), a 24-hour (*N* = 45; 15 3-year-olds, 15 4-year-olds, and 15 5-years-olds), or a 1-week (*N* = 47; 15 3-year-olds, 15 4-year-olds, and 17 5-years-olds) delay.

As shown in Figure 1, during the encoding phase, child participants were presented with eight unique trials of animal-scene associative pairs in a timed sequence. Within each association trial, an interactive storybook showed children an isolated animal image in full screen, then a scene image as the animal’s “favorite place” on the next screen, followed by a third screen showing an image that placed the animal within the same scene image. Each screen was presented for 5 seconds with speech narration. The pool of images consisted of eight animals and 20 scenes. There were four storybook versions in total, and each contained the same eight animals paired with eight different scenes. Children were randomly assigned to one of the four storybook versions (Storybook 1: *N* = 37; Storybook 2: *N* = 34; Storybook 3: *N* = 33; Storybook 4: *N* = 33). During the memory test phase (after 5 minutes, 24 hours, or 1 week depending on condition), child participants completed eight 4AFC (alternative forced-choice) trials with speech narration. Within each test trial, child participants were shown one of the eight animal images (*cue*) on a screen for 5 seconds, followed by a screen for an unlimited time with the cue animal in the center and four scene options surrounding the animal. The four scene options consisted of the correct scene (*target*) associated with the animal during encoding, two lure scenes associated with other animals during encoding, and a novel scene missing from the encoding phase. All scene choices came from the pool of 20 scene images. Child participants did not receive feedback on whether they chose the correct scene. Saragosa-Harris et al. (2021) observed that children’s memory accuracy increased with age and decreased with longer delays.

**Figure 1.**
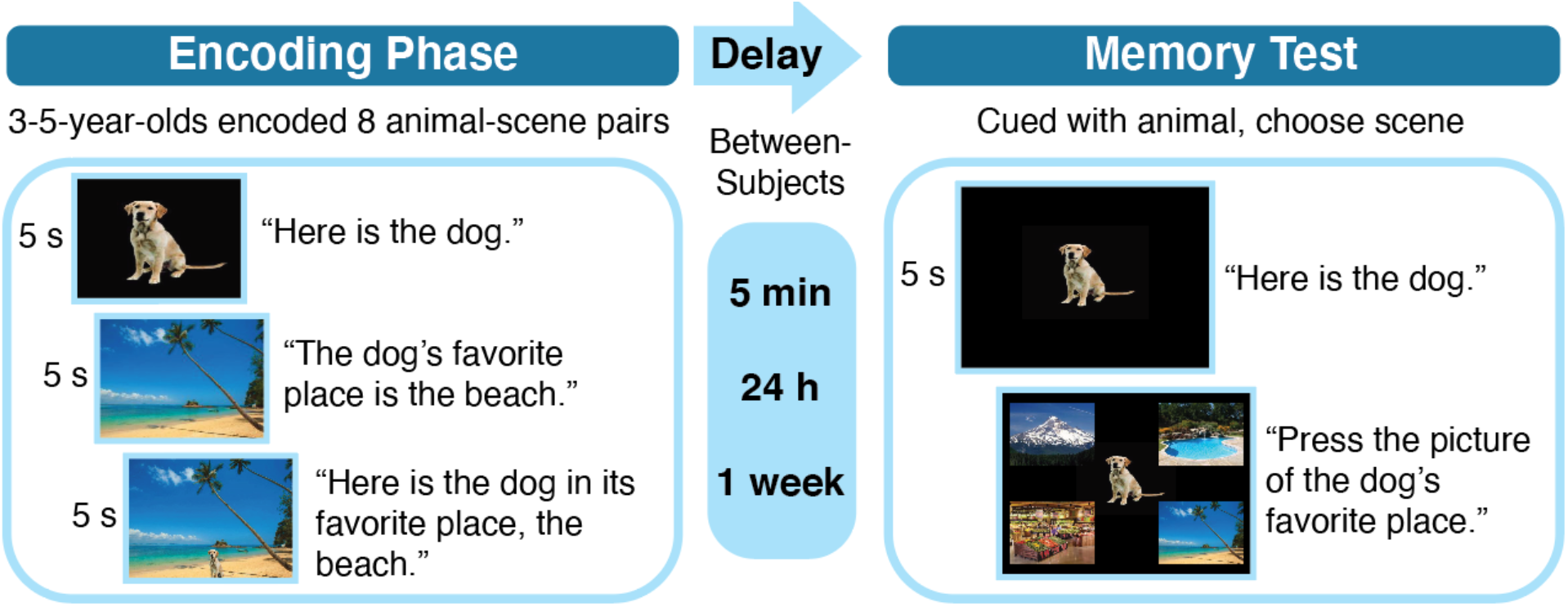
Procedure to obtain children’s memory. *Note*. During the encoding phase (left panel), a speech narration described for child participants each animal, then its favorite place, followed by the animal in the place. Each screen was presented for 5 s. Each child participant encoded eight animal-scene pairs. After a 5-min, 24-h, or 1-week delay (between subjects), child participants took a memory test (right panel) consisting of eight trials. Within each trial, participants first viewed an animal *cue* for 5 s with narration and were then asked to choose the correct associated scene (*target*) out of four scenes.

### ResMem Memorability Score Predictions

We inputted the 59 target scenes and the 60 target animals from the adult experiment (see next section), including images from the child dataset, into ResMem and extracted a predicted memorability score (*hit rate*) for each image. ResMem is a pre-trained residual deep neural network (DNN), which combines AlexNet feature layers that capture low-level visual information with ResNet feature layers that capture high-level semantic information of the inputted images (Needell & Bainbridge, 2022). ResMem was trained to predict image memorability using large-scale adult memory performance data obtained in similar continuous recognition tasks as the adult memory experiment in this study. The predicted memorability score ResMem assigns to each image, ranging from 0 to 1, indicates the probability that this image is successfully recognized among adults when repeated. Images with higher predicted memorability scores are better remembered by adults (Spearman correlation *rho* = 0.67; Needell & Bainbridge, 2022). Hence, ResMem predictions can be treated as a proxy for adults’ memory.

### Adult Memory Experiment

To confirm that the memorability scores predicted by ResMem DNN are a reliable proxy for adults’ memory performance, we ran two continuous recognition experiments among adults and calculated the memorability score (*hit rates*) for each of the 59 scene and 60 animal images. Note that while children’s memory was measured with an associative memory task, here adult memory is measured with an item-based continuous recognition task. Prior work has shown that memorability measures are stable across different task parameters and image contexts, and can be measured using both item-based and associative tasks (Bainbridge, 2020; Bylinskii et al., 2015; Xie et al., 2020). However, we revisit the differences between these two tasks in the Discussion.

#### Participants

A hundred and sixteen online participants on Amazon Mechanical Turk engaged in a scene memory experiment, where a timed sequence of scene images was shown, and 148 online participants engaged in an animal memory experiment, where a timed sequence of animal images was shown. Prior work showed that having 80 participants in a continuous recognition task was optimal for estimating memorability scores for each image (Isola et al., 2014). We recruited 160 online participants for each experiment to account for potential exclusions. After applying the exclusion criteria (see the end of the Procedure subsection), 116 participants from the scene experiment and 148 participants from the animal experiment were included for analysis. Participants consented to their participation following the guidelines of the Institutional Review Board (IRB) at the University of Chicago (IRB19-1395). Participants could participate in both experiments, and each participant was awarded a $0.70 compensation upon completion. Each experiment took approximately seven minutes, and participants were allowed to end the experiment at any time.

#### Stimuli

A pool of 136 scene images was shown in the scene experiment, including the 20 scene images from the child dataset and 116 from the Scene Understanding (SUN) database (Xiao et al., 2010). From the pool of 136 scene images, 76 were used as *filler* images, and 60 were used as *target* images, including the 20 from the child dataset. We included more images in the adult continuous recognition tasks beyond the 20 images from the child dataset in order to increase task difficulty. The higher number of images allows for a larger time interval between target encoding and recognition, and also ensures enough statistical power during image-based analyses. We balanced the proportions of indoor, natural outdoor, and manmade outdoor images between the target and the filler sets. To avoid images coming from the child dataset versus from the SUN database being visually different and thus leading to systematic variations in visual memory, we prioritized selecting scenes from the SUN database which contained themes (e.g., forest, sea, house) that were similar to the 20 scenes from the child dataset. For both experiments, we ensured that the image set from the child dataset versus the one we adapted online did not differ in low-level visual features using the Natural Image Statistical Toolbox (Bainbridge & Oliva, 2015) (all *p* > .05). All images were resized to 256 × 256 pixels. One target scene image from the SUN database was misaligned in dimensions with the rest of the scene images and was excluded from analysis in the Results section, resulting in 59 scene images that we used for analyses.

A pool of 136 animal images was shown in the animal experiment, including the eight animal images from the child dataset and 128 from Google Images (with Creative Commons licenses). To ensure that the two sets of animals were visually similar, we selected Google images that contained isolated animals with neutral poses and expressions. We photoshopped the Google images following the format of the eight animal images from the child dataset by placing each isolated animal in the middle of a black background. The 136 animals came from different species that are common in American zoos and aquariums. Similar to the scene experiment, 76 animal images were used as fillers, and 60 were used as targets, including the eight from the child dataset. We balanced the proportions of mammals, fish, reptiles, insects, and birds between the target and the filler sets. All images were resized to 256 × 256 pixels.

#### Procedure

After answering three common-sense questions at the beginning of each experiment (e.g., “What is the name of the last month of the year?” and “What is the capital of the U.S.?”), participants were directed to the main task. The main task adopted a continuous recognition paradigm frequently used to test image memorability (Bainbridge et al., 2013), where participants were shown a timed sequence of images and were instructed to press the *r* key when they recognized an image from earlier in the sequence (Figure 2). Each image was presented for 750 ms with an 800-ms interstimulus interval where participants looked at a fixation cross. Each of the 136 images within each experiment was repeated at most once. All target images, but not all filler images, were repeated. Target repetitions were spaced at least 30 seconds apart. Filler repetitions were spaced 0-4 images apart and were included to maintain participant attention.

**Figure 2.**
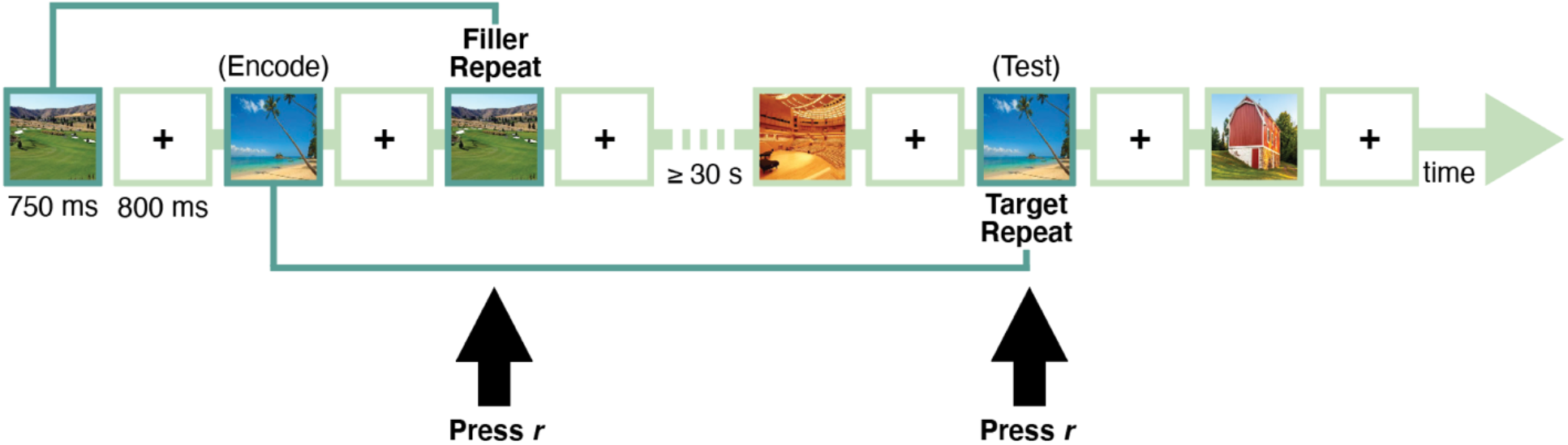
A flowchart of the adult visual memory experiment for scene images. *Note*. Each image was presented for 750 ms, followed by an 800-ms fixation cross. Participants pressed *r* when they saw an image repeat. Memorability scores were derived for target images, which were repeated at least 30 seconds apart. Filler repetitions were spaced 0-4 images (less than 6 seconds) apart. The adult experiment for animal images followed the same paradigm.

To ensure the quality of participants, only workers with an AMT approval rate of at least 95% and the number of approved tasks of at least 50 were allowed to participate. Only workers with an IP address within the United States were allowed to participate. Participants who answered any of the three common-sense questions incorrectly were excluded. Participants who paid no attention and indicated repetitions for over 95% of the images in the timed sequence (i.e., kept pressing the *r* key) were excluded. We also used filler repetitions as a vigilance test to exclude participants who paid little attention during the experiments. Participants with over 70% false alarms on the first presentation of filler images and over 70% misses on vigilance repeats were excluded.

### Analyses

#### Calculating Memory Performance

To compare children’s and adults’ memory, we generated item-wise memory measures between 0 and 1 for target images from the child dataset and the adult experiment. We obtained children’s memory accuracy for each of the 20 scene images by calculating the percentage of trials a scene was chosen when it was the target scene (*percent correct*). We obtained children’s memory accuracy for each of the eight animal images by calculating the percentage of trials the target scene was chosen when cued with an animal. To obtain adults’ memory accuracy, we measured the adult experiment hit rates (*adult HR*) on image repetitions of the 59 target scene images and the 60 target animal images which included images from the child dataset. Finally, we also obtained predictions of memorability scores (hit rates) for each of the images through the ResMem DNN (*ResMem HR*). Figure 3 shows examples of scene and animal images that are memorable and forgettable according to memory scores extracted from ResMem, the child dataset, and the adults’ experiments.

**Figure 3.**
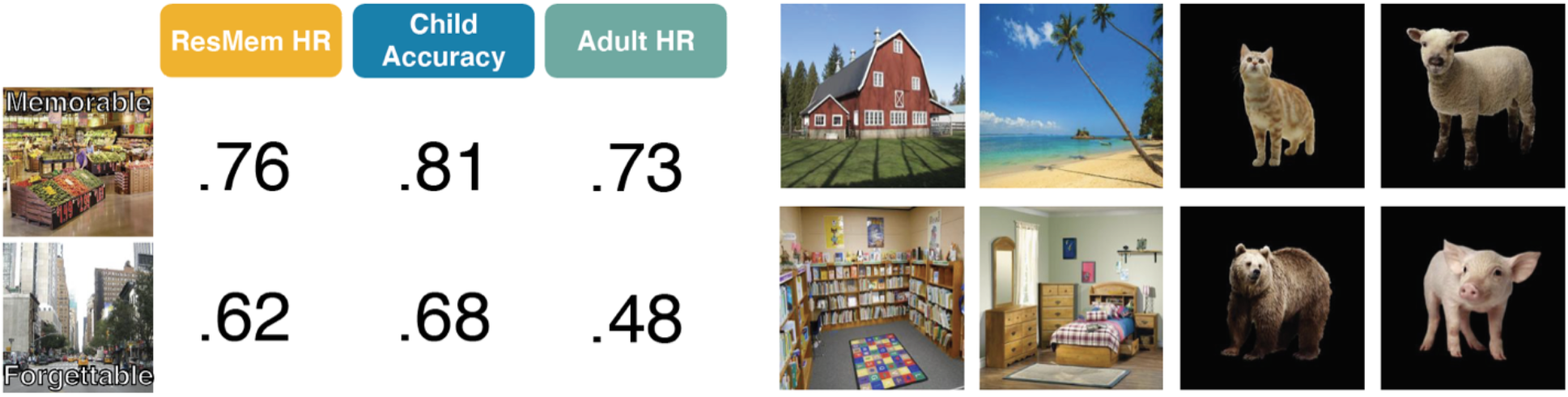
Example images from the child dataset and memory measures. *Note*. Left: Examples of memorable and forgettable scene images, marked with single-score measures of ResMem-predicted memorability (*ResMem HR*), children’s memory accuracy, and adults’ memory (*adult HR*) for each image. Right: More examples of scene and animal images from the child memory dataset.

#### Logistic Regression Model

We generated a logistic regression model to predict children’s memory accuracy using trial-wise test data from the child dataset, where each child answered eight test trials. Logistic regression was chosen because trial outcomes were binary (i.e., correct vs. incorrect answers). Age group (3-year-olds, 4-year-olds, and 5-year-olds) and the three delay conditions (5-min, 24-h, and 1-week) were entered as categorical variables into the regression model. We z-transformed continuous data before fitting the model. The model included ResMem-predicted memorability score (*ResMem HR*) of the target scene in each test trial, *age*, and *delay* as predictors:

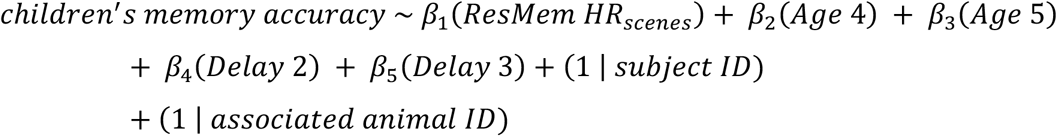

Note that we included two random intercepts. The random intercept of child participant ID accounted for individual differences across children. The random intercept of the associated animal image ID (i.e., 1-8) in each test trial accounted for the fact that child participants encoded scene images in association with animal images. The rationale was that even if different children encoded an animal associated with different scenes, the same animal might affect children’s memories of different scenes in a similar way. We call this model the base model (see Supplemental Materials for other models we tested; this base model had the best fit metrics adjusted R^2^ and AIC of all models).

### Constraints on Generality

We used an online sample representative of the U.S. population to collect adults’ behavioral data. ResMem was also trained using adults’ behavioral data from an online sample representative of the U.S. population. However, these online participants were adults in the United States with internet access, which may limit the generalization of results in this study to all human adults, even though memorability has been shown to be robust across populations. In addition, the original child dataset came from a New York area sample, which could limit the generalizability of the results.

### Transparency and Openness

We report how we determined our sample size, all data exclusion criteria, all manipulations, and all measures in the study. All data are available at https://osf.io/pz26c/ (data will be made available upon publication of the paper). Data were analyzed using MATLAB, version 9.11.0.1769968 (R2021b). This study’s design and its analysis were not pre-registered.

## Results

We have generated single-score measures of children’s memory (percent correct), ResMem-predicted memorability (ResMem HR), and adults’ memory (adult HR) for all target images. To find out how early children have developed adult-like visual memory patterns, we compared children’s memory accuracy with image memorability extracted from ResMem and the visual memory behavioral experiment (adult experiment). Because images with higher hit rates are more likely to be remembered consistently across adults (Bainbridge et al., 2013; Isola et al., 2011), we could use whether images with higher adult HR were also better remembered by children as an indicator of whether children have developed adult-like visual memory by a certain age. As an overview, we first compared ResMem HR with adult HR on the same set of images to ensure that ResMem predictions successfully captured the image memorability effects on adults. We then compared children’s memory accuracy to ResMem HR and adult HR, respectively, on the same set of images to see whether ResMem predictions and adults’ memory could predict children’s image memory.

### Does ResMem predict adults’ memory?

As a sanity check for the adults’ memory data, we tested the reliability of adult HR for the 59 scene targets (*M* = .65, *SD* = .07, min = .48, max = .74) by splitting adults into random halves and running a Spearman rank correlation between the 59 memorability scores from each half, as shown in Figure 4A. A higher similarity between the two score ranks would indicate higher consistency in adults’ memory. Across 1000 of these random split-half trials, the averaged Spearman-Brown corrected *rho* was .73 (*p* < .001, calculated using a permutation test with 1000 shuffled rankings). Figure 4B shows results of the same split-half consistency analysis conducted using the 60 animal targets (*M* = .73, *SD* = .04, min = .64, max = .76, averaged Spearman-Brown corrected *rho* = .79, *p* < .001). These results showed that adults were indeed highly consistent in what scene and animal images they remembered and forgot.

**Figure 4.**
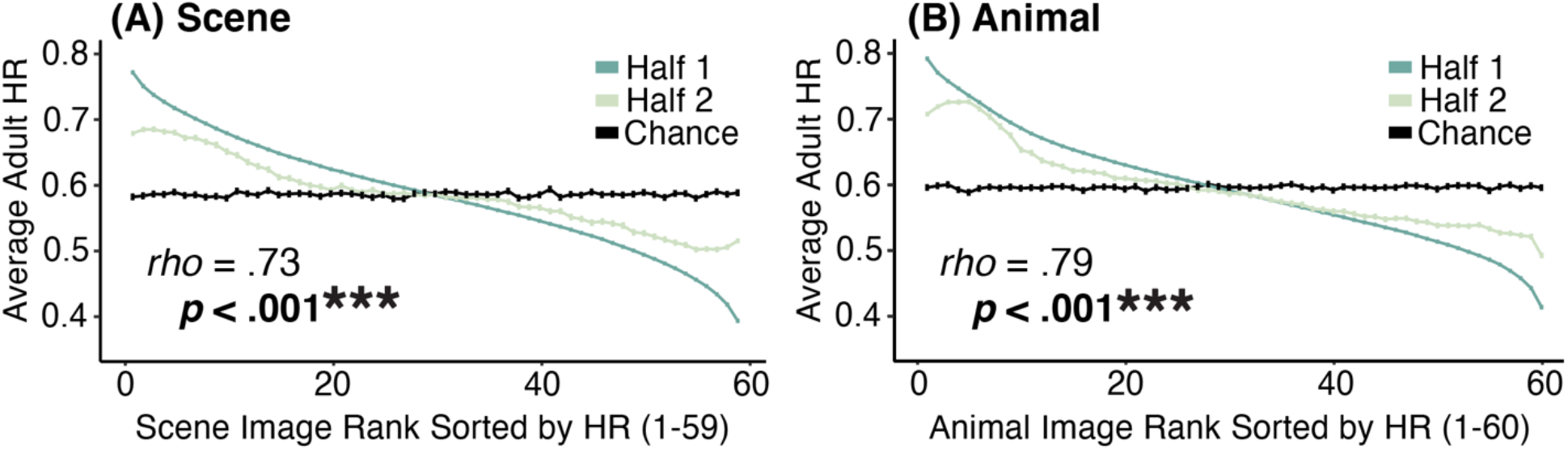
Consistency graphs for adult hit rate (adult HR) *Note*. (A): Consistency graph for adult hit rate (adult HR) of the 59 scene images. (B): Consistency graph for adult hit rate (adult HR) of the 60 animal images. Adult participants were split into random halves in each of the 1000 iterations. The memorability scores calculated from the first half of the participants (Half 1: dark green line) were ranked from high to low, and those from the second half (Half 2: light green line) were ranked using the image rank of Half 1. The two green lines represent the average of adult HR ranks over the 1000 iterations. If the two halves of participants are perfectly consistent in memory, the green lines should coincide. In contrast, if the two halves of participants are not consistent in memory, the light green line should coincide with the chance line in black, which represented a random shuffle of image ranks in Half 1. The significant Spearman rank correlation indicates that split halves of participants tend to remember and forget the same images. Error bars represent standard error of the mean.

We then tested whether ResMem predicted higher memorability scores for images that were better remembered by adults, given that ResMem was trained to predict image memorability using large-scale adult’s memory data on photographs. Such a sanity check provides a benchmark of whether ResMem predictions can be treated as a proxy of adults’ memory. To see whether ResMem (*M* = .73, *SD* = .11, min = .50, max = .92) successfully assigned higher memorability scores to more memorable scene images, we ran a Spearman correlation between the 59 memorability scores extracted from ResMem and those calculated from adults’ memory data, as shown in Figure 5A. As expected, ResMem HR significantly correlated with adult HR for the scene images (*rho* = .54, *p* < .001). This result means that ResMem HR can be treated as a reliable proxy of adults’ memory of the scene images in further analyses. Figure 5B shows results of the same Spearman correlation performed over the 60 animal targets. Results showed that ResMem HR (*M* = .86, *SD* = .05, min = .80, max = .95) did not correlate with adult HR of animal images (*rho* = .22, *p* = .90), meaning that ResMem HR was not a good substitution for adults’ memory of animal images. We revisit ResMem’s performance with the animal images in the Discussion. Because ResMem HR could not be treated as a reliable proxy of adults’ memory for the animal images, we will mainly focus on results for scene images.

**Figure 5.**
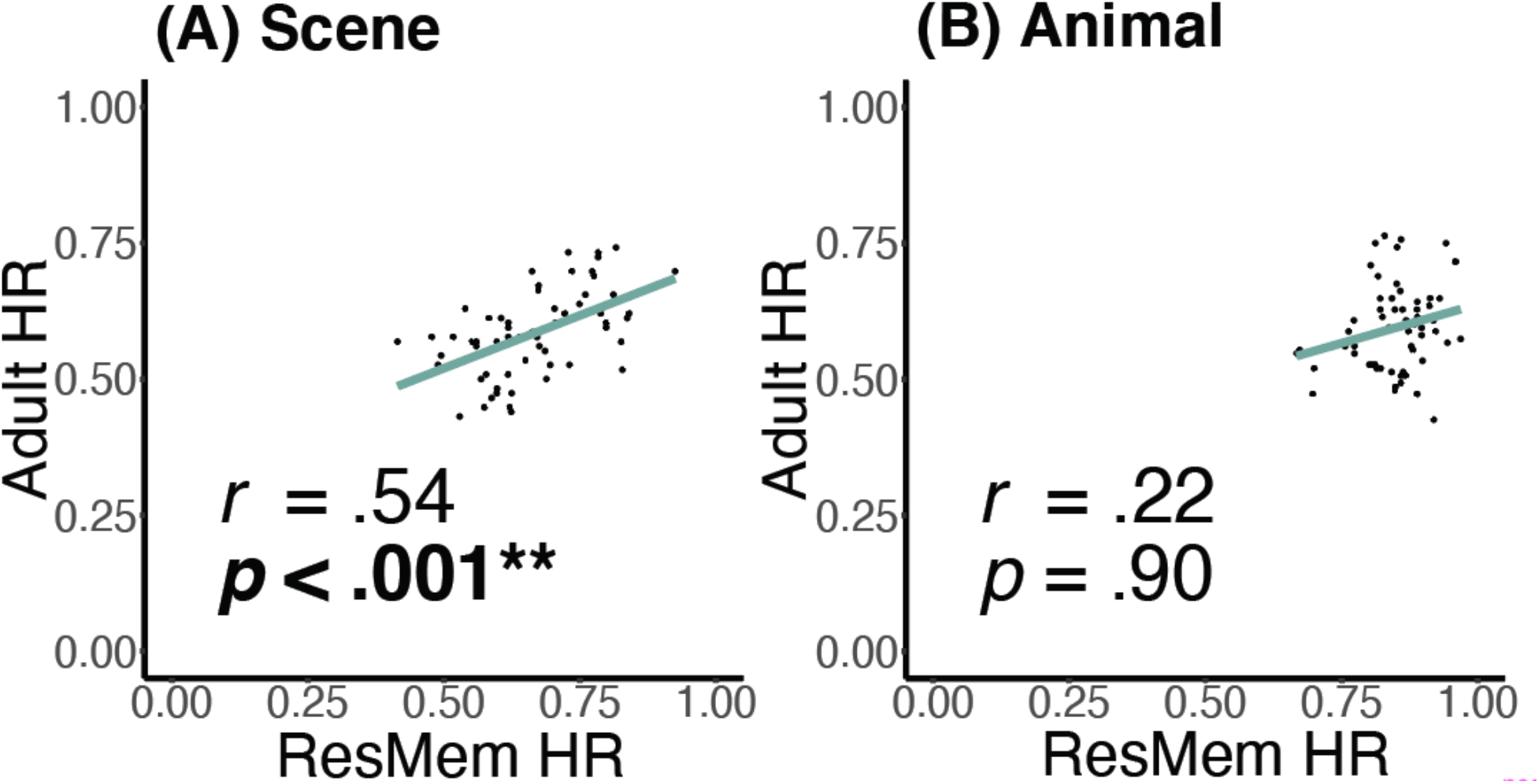
Spearman correlation graphs of ResMem HR vs. adult HR. *Note*. (A): Spearman correlation graph of ResMem HR vs. adult HR on the 59 scene images. ResMem HR significantly predicted adult HR on the 59 scene images, meaning that ResMem HR is a good proxy of adults’ memory of the scene images. (B): Spearman correlation graph of ResMem HR vs. adult HR on the 60 animal images. ResMem HR did *not* predict adult HR on the 60 animal images, meaning that ResMem HR is *not* a good proxy of adults’ memory of the animal images.

### Does ResMem predict children’s memory?

The central question of this study is by what age children have shown adult-like visual memory patterns. To see which child age group remembered memorable images more than forgettable ones, we Spearman correlated ResMem-predicted memorability with children’s memory accuracy within each age group across delay conditions (3-year-olds: *M* = .52, *SD* = .19; 4-year-olds: *M* = .62, *SD* = .15; 5-year-olds: *M* = .76, *SD* = .07) using the 20 scene images from the child dataset. Figure 6A-C shows the crucial results: ResMem HR did not predict children’s image memory at the age of 3 (*rho* = -.15, *p* = .53), but significantly predicted children’s image memory at the ages of 4 (*rho* = .52, *p* = .02) and 5 (*rho* = .58, *p* = .007) (FDR-corrected, *q* < 0.05). This means that children develop adult-like sensitivity to image memorability by the age of 4. To see whether different delay intervals between encoding and memory test phases affected the image memorability effect on children’s memory, we Spearman correlated ResMem HR with children’s memory within each delay condition across age groups (5-min delay: *M* = .79, *SD* = .09; 24-h delay: *M* = .66, *SD* = .13; 1-week delay: *M* = .47, *SD* = .11) using the same 20 scenes. ResMem HR did not predict children’s image memory across age after a 5-minute (*rho* = .29, *p* = .21) or a 24-hour delay (*rho* = -.15, *p* = .52), but significantly predicted children’s image memory after a 1-week delay (*rho* = .55, *p* = .01) (FDR-corrected, *q* < 0.05), as shown in Figure 6D-F. We confirmed these results by combining 4- and 5-year-olds into one group and running the same Spearman correlation within each delay condition. We found the same results, where ResMem HR predicted children’s memory after a 1-week delay (*rho* = .65, *p* = .002) (FDR-corrected, *q* < 0.05) but not after shorter delays (*ps* > .05). This means that the memorability effect drives children’s image memory accuracy after long delays.

**Figure 6.**
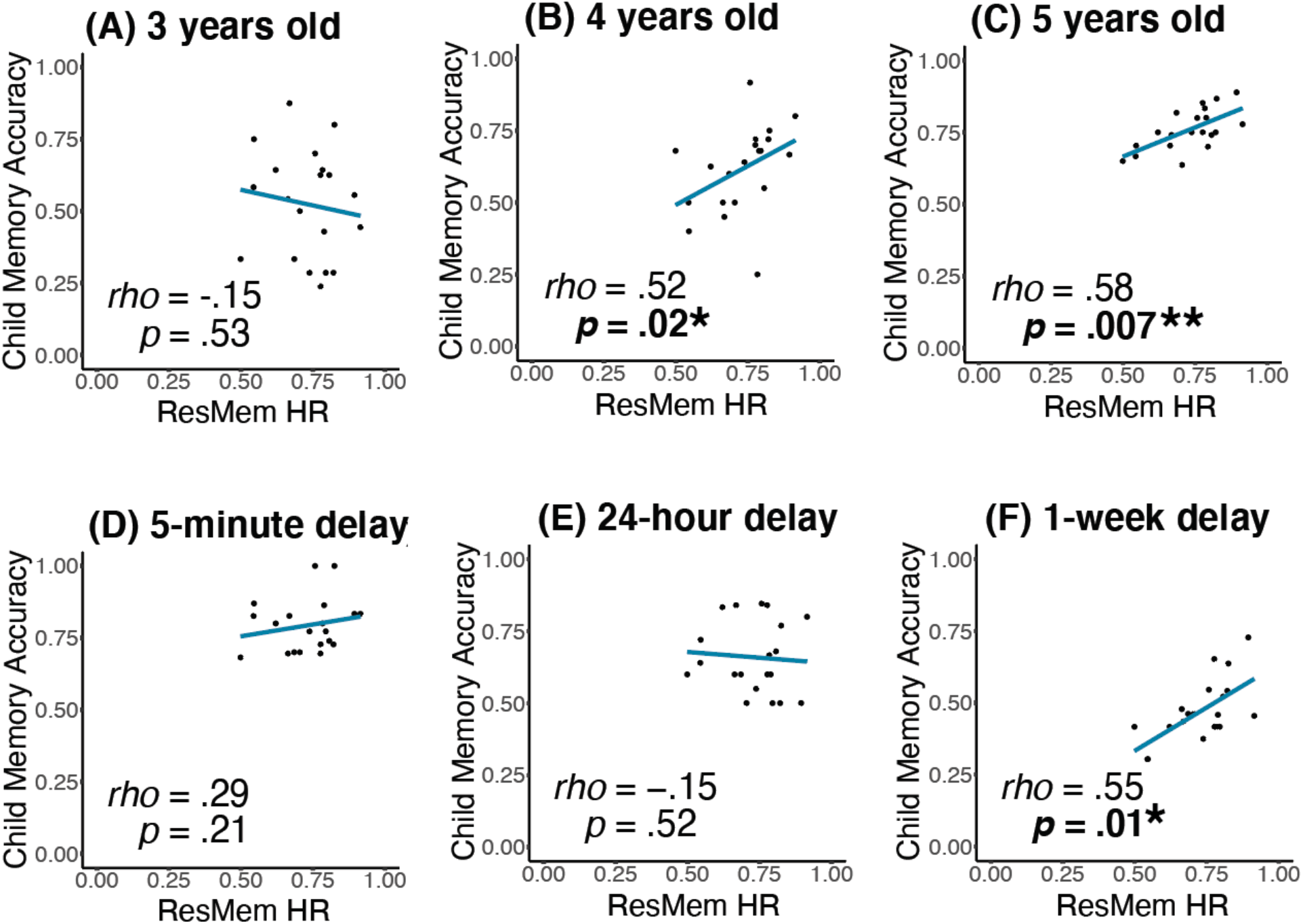
Spearman correlation graphs of ResMem HR vs. children’s memory accuracy. *Note*. Top (A-C): Spearman correlation graphs of ResMem HR vs. children’s memory accuracy in each age group across delay conditions on the 20 scene images from the child memory dataset. (A) ResMem did not predict the memory of 3-year-olds across delays. (B, C) ResMem significantly predicted the memory of 4- and 5-year-olds across delays. Bottom (D-F): Spearman correction graphs of ResMem HR vs. children’s memory accuracy in each delay condition across age groups on the 20 scene images from the child memory dataset. (D, E) ResMem did not predict children’s memory across age after a 5-minute or a 24-hour delay. (F) ResMem significantly predicted children’s memory across age after a 1-week delay.

Knowing that both age and delay could influence how well ResMem predicted children’s image memory, we wanted to create a statistical model that could predict children’s memory using ResMem HR by specifying children’s ages and delay intervals. We fitted the 1096 test trials to our base regression model (see Method), which included ResMem HR predictions for scenes, child’s age, delay, and random intercepts of subject ID and associated animal ID, and found a significant model fit (adjusted *R^2^* = .39, *p* < .001, AIC = 5069). As expected, we found that the predictor of ResMem HR for scenes was significant (*β_ResMem HR Scenes_* = .85, *p* = .01). We also found that both age predictors were significant (*β_Age 4_* = .70, *p* = .02; *β*_Age *5*_ = 1.31, *p* < .001). Both delay predictors were also significant (*β_Delay 2_* = -.94, *p* = .002; *β_Delay 3_* = −1.95, *p* < .001). This means that the intrinsic memorability of scenes, children’s age, and time delay, all had a significant effect on children’s memory. While we tested other model structures, this base model led to the best model fit (see Supplemental Materials for statistics of all models).

### Possible explanations for the change in children’s memory

One potential hypothesis to account for the fact that ResMem could predict the scene memory of 4- and 5-year-olds but not that of 3-year-olds is that 3-year-olds’ memory might be noisier and less consistent across participants compared to older children’s, given that their memory performance is generally worse (Saragosa-Harris et al., 2021). To test this hypothesis, we ran a split-half consistency analysis for each age group using the 20 scene images. Similar to the adult consistency analyses mentioned earlier, we split children within each age group into random halves and ran a Spearman correlation between the 20 data points from each half over 1000 iterations, as shown in Figure 7. To our surprise, 3-year-olds were consistent in the scene images they remembered and forgot (averaged Spearman-Brown corrected *rho* = .65, *p* = .007), whereas 4- (averaged Spearman-Brown corrected *rho* = .40, *p* = .14) and 5-year-olds (averaged Spearman-Brown corrected *rho* = -.72, *p* = .85) were not consistent in memory. We acknowledge that the nonsignificant results for the two older groups might be due to low power considering the low number of participants within each age group (45 3-year-olds, 45 4-year-olds, and 47 5-year-olds) as well as the low number of images (20; see Supplemental Materials for more results on this point). Nevertheless, the reliable memory pattern at the age of 3 ruled out the hypothesis that young children might have noisier, inconsistent memory patterns. Given that ResMem could not predict the scene memory of 3-year-olds, these results suggest that 3-year-olds might consistently use a common encoding strategy that ResMem does not capture. We tested different possible strategies in a Supplemental Experiment S1 (see Supplemental Materials), where adults rated how much 3-year-olds would be familiar with or attracted to each scene image. For example, the most familiar scene images for 3-year-olds were those of a playground, bounce house, and library. Familiarity (*rho* = .49, *p* = .03) but not aesthetic rating (*p* > .05) of the scene images could predict the memory of 3-year-olds, suggesting that image familiarity, but not aesthetics, may influence memory at the age of 3.

**Figure 7.**
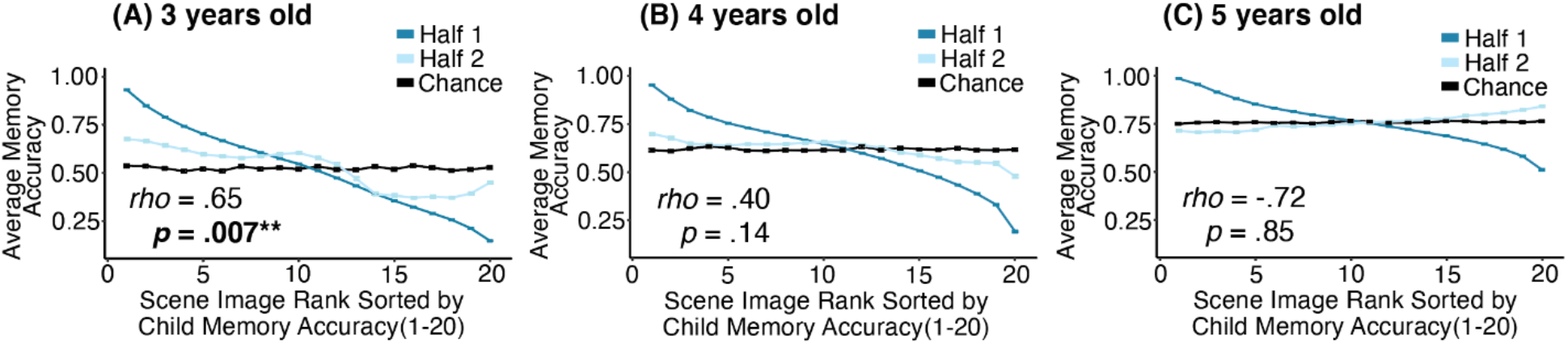
Consistency graphs for children’s memory accuracy within age. *Note*. Within-age consistency graphs for children’s memory accuracy of the 20 scene images from the child memory dataset. Child participants within each age group (*N* ≈ 45) were split into random halves in each of the 1000 iterations. The 20 data points (image memory accuracy scores) from the first half of the children (Half 1: dark blue line) were ranked from high to low accuracy, and the 20 data points from the second half (Half 2: light blue line) were ranked using the image rank of Half 1. The two blue lines represent the average accuracy at each rank over the 1000 iterations. If the two halves of children are perfectly consistent in memory, the blue lines should coincide. In contrast, if the two halves of participants are not consistent in memory, the light blue line should coincide with the chance line in black, which represented a random shuffle of image ranks in Half 1. (A): 3-year-olds consistently remembered and forgot certain scene images. (B,C): 4- and 5-year-olds did not show consistent memory patterns of scene images within each age group. Error bars represent standard error of the mean.

An additional explanation is that children might begin to develop adult-like encoding strategies around the age of 4. Given that 3-year-olds had consistent memory patterns that could not be predicted by ResMem, whereas 4- and 5-year-olds had inconsistent memory patterns that could be predicted by ResMem, we conjectured that the inconsistent memory pattern at ages 4 and 5 might indicate a mix of encoding strategies. Specifically, some 4- and 5-year-olds might have developed adult-like strategies, while others remained using the same strategies as 3-year-olds. Because age is continuous and development takes place gradually, older 4-year-olds may show more similar memory patterns to adults than younger 4-year-olds. If this is the case, then we would expect to see ResMem better predicts the memory of older 4-year-olds than that of younger 4-year-olds. To test this hypothesis, we split 4-year-olds into a younger sub-group (age = [4, 4.5), *N* = 18, *M* = .69, *SD* = .17) and an older sub-group (age = [4.5, 5), *N* = 27, *M* = .55, *SD* = .20), and Spearman correlated ResMem HR with children’s memory accuracy within each sub-group using the 20 scene images from the child dataset. As expected, ResMem HR did not predict the memory of the younger 4-year-old sub-group (*rho* = .34, *p* = .14), but significantly predicted that of the older sub-group (*rho* = .48, *p* = .03) (Figure 8). These results suggest that the emergence of adults’ memory patterns in early childhood is likely accompanied by the development of adult-like visual encoding strategies.

**Figure 8.**
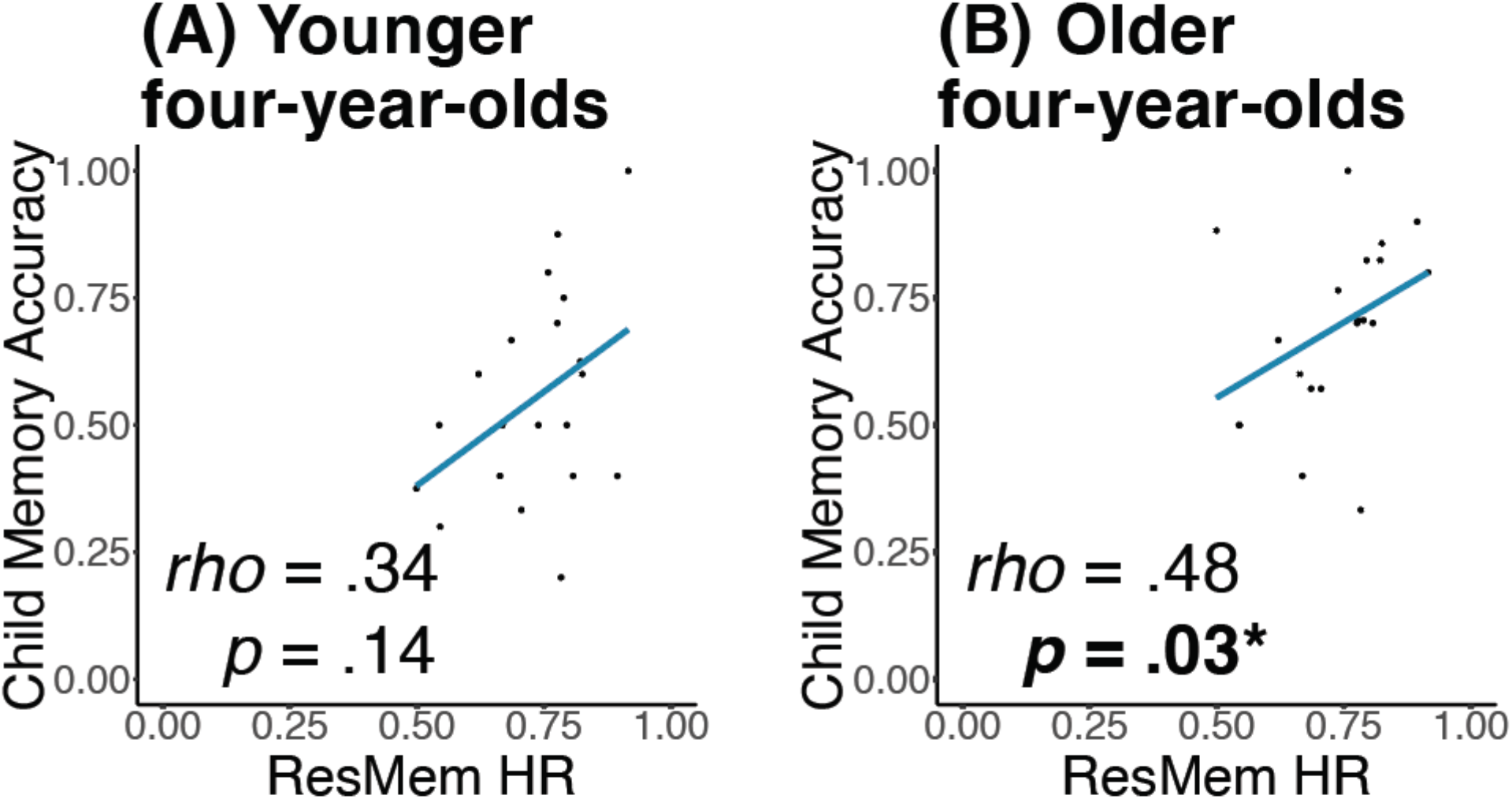
Spearman correlation graphs of ResMem HR vs. children’s memory accuracy at the age of 4. *Note*. Spearman correlation graphs of ResMem HR vs. children’s memory accuracy at the age of 4 on the 20 scene images from the child memory dataset. The 4-year-olds (*N* = 45) were separated into a younger and an older group by month. (A): ResMem did not predict the memory of the younger 4-year-old group (4-4.5 years). (B): ResMem significantly predicted the memory of the older 4-year-old group (4.5-5 years).

### Does adults’ memory predict children’s memory?

Knowing that ResMem predicted children’s memory in older age groups and after a long delay, we expected adults’ memory to predict children’s memory, because ResMem HR was a reliable proxy of adults’ memory. We ran Spearman correlations between adult HR and children’s memory accuracy within each age group and delay condition using the 20 scene images from the child dataset. None of the correlations yielded significant results (all *p* > .05, see Supplemental Materials for detailed statistics), highlighting that ResMem outperformed adults’ memory data in predicting children’s memory. A potential explanation is that adult behavioral data was prone to sampling error. Adult HR (*M* = .65, *SD* = .07) had lower variation compared to ResMem HR (*M* = .73, *SD* = .11), meaning that the differences in memorability may be less clear across these specific images in adults.

To see whether ResMem HR and adult HR predicted children’s memory of animal images, we ran the same Spearman correlations within each age group and delay condition using the eight animal images. None of the correlations yielded significant results (all *p* > .05). Such nonsignificant results might-be because there were only eight data points in each correlation due to a low number of animal images. Children also had to remember a combination of the animal cues and the scene targets in order to perform the associative memory task. Hence, another potential explanation is that children’s memory is less driven by the visual features of the cue items than the target items during this type of task. See more discussion on this point in the General Discussion.

## Discussion

Here, we discover that children develop adult-like visual memory patterns by the age of 4, and ResMem predictions of adult memorability were able to predict children’s memory for scene images, particularly after a 1-week delay. Unexpectedly, children at the age of 3 showed consistent memory patterns within their age group and likely remembered images depicting familiar scenes, whereas older children stop showing consistent memory patterns within each age group, reflecting a period of visual memory development of becoming more adult-like. We have also created a novel logistic regression model that could be used to predict children’s scene memory accuracy using ResMem predictions, age, and delay.

Note that although children showed adult-like visual memory by the age of 4, it does not mean that children younger than 4 remembered little. Interestingly, children may apply different visual memory strategies from adults before they adopt adults’ encoding strategies. We found that 3-year-olds did not show adult-like memory patterns but consistently remembered scenes that were rated as familiar to children at the age of 3. In contrast, 4- and 5-year-olds exhibit adult-like memory patterns despite not showing consistent memory within each age group. This means that 3-year-olds reliably remember certain images, and 4- and 5-year-olds start to develop adult-like visual memory strategies. We discuss potential mechanisms underlying such development in the next subsection (*Potential mechanisms for the development of adult-like memory patterns*).

There are a few possible explanations for why ResMem could predict children’s memory after a long, 1-week delay but not after shorter delays. One possibility is that the information used by ResMem to predict memory better matches the information remaining in memory after a delay. Specifically, given the decay of visual information in memory over time observed in adults (Lampinen et al., 2001; Potter, 2012), it is possible that the information that persists in memory is what most drives memorability effects. For example, richer semantic features of an image have been shown to be most predictive of memorability (Kramer et al., 2022) and could be the information that remains in memory after a delay (Lampinen et al., 2001; Potter, 2012). It is also possible that other factors such as attention play a stronger role than memorability at shorter delays. Although the memorability effect remains stable with different delays among adults (Isola et al., 2014, p. 201; Lin et al., 2021), it is unclear whether it pertains across time among children. Finally, another possible explanation is that there might have been a ceiling effect in children’s memory after shorter delays, especially after the 5-minute delay, making it difficult for ResMem HR to predict children’s memory performance.

### Potential mechanisms for the development of adult-like memory patterns

We raised two competing hypotheses of the emergence of humans’ sensitivity to the memorability effect. One hypothesis is that infants develop such sensitivity through neural maturation or visual information accumulation (the developmental account). Through development, infants’ visual memory emerges from non-adult-like to adult-like between ages 3 and 4. Another hypothesis is that from very early on without any postnatal developments, infants possess sensitivity to the memorability effect and show adult-like visual memory patterns. Results from the current study support the developmental account. Children’s memory patterns develop to be more adult-like and become predictive by ResMem, a proxy of adults’ memory, by age 4. Another piece of evidence is that older 4-year-olds showed more adult-like memory patterns compared to younger 4-year-olds. This implies that the memory of older children was more adult-like compared to their younger counterparts both within and across age groups.

One possible mechanism behind the developmental account lies in neural changes in early childhood. We have noted that a decline in neurogenesis in the hippocampus improves its functional competence in holding long-term memory and leads to the fading of infantile amnesia by the age of 4 (Akers et al., 2014; Bauer et al., 2011; Dennis et al., 2016; Schneider & Pressley, 2013). Results from this study suggest a temporal coincidence between the disappearance of infantile amnesia and the emergence of adult-like visual sensitivity. Previous research suggests a role of the medial temporal lobe and hippocampus in sensitivity to memorability by prioritizing memorable perceptual information during encoding (Bainbridge et al., 2017; Bainbridge & Rissman, 2018). We thus conjecture that the maturity of the hippocampus through a neurogenesis decline may have also supported adult-like visual sensitivity at around age 4. Additionally, a neural change in plasticity across development (Scher, 2008; Scher & Loparo, 2009) may also explain the emergence of adult-like visual memory. Children up to the age of eight can recognize faces from all races equally well, whereas older children and adults recognize own-race faces better than other-race faces (Goodman et al., 2007). With a decline in neural plasticity, the fluidity in visual perception declines, which shapes the perceptual expertise in one category of face stimuli. Similarly, young children may have high fluidity and are sensitive to diverse visual statistics during image encoding, whereas adults are sensitive to certain visual statistics that constitute the memorability effect, shaped by a decline in neural plasticity. It is possible that children have an established sense of certain visual statistics by age 4 and thus tend to remember the same images as adults.

Another potential mechanism underlying the developmental account may be the accumulation of cognitive experience. It is possible that by the age of 4, children have accumulated enough visual samples from the world to support adult-like visual processing or an adult-like model of visual memory. Around the age of 4, children start to attend kindergarten and engage in more diverse experiences, which may also result in children acquiring more adult-like visual experiences. In addition, the accumulation of language inputs leads to a rapid improvement in young children’s language skills, which may allow children to extract high-level information and encode abstract representations of scene images by 4 years of age. Abstract, semantic information may be the type of features that predominantly drive consistencies in memorability (Kramer et al., 2022; Needell & Bainbridge, 2022).

### Associative and item-based measures of memorability

In this study, we compared children’s memory obtained from an associative memory task with adult memory and neural network predictions obtained from an item-based recognition memory task. Child participants encoded animal-scene image associations and were tested on scene recognition with animal cues after delays. The adult experiment did not have a separate memory test phase; instead, the continuous recognition paradigm showed repeated targets in a timed sequence and asked adult participants to indicate target recognition. The ResMem DNN was trained on data with different images from the same continuous recognition task paradigm. Despite these task differences, ResMem is remarkably able to predict children’s memory for targets in an associative memory task. In fact, Xie et al. (2020) showed that in an associative word recall task, the memorability of the target word better predicted participants’ recall memory performance than that of the associated word cue. Indeed, the logistic regression results in this study show that the memorability of the target scenes rather than animal cues predicts children’s memory accuracy. This provides evidence that the memorability of the target items being recalled drives memory success in an associative memory task.

It remains a question why ResMem does not predict children’s and adults’ memories of animal images. One possible explanation is that ResMem was trained using natural images, whereas the animal images used in this study might have become unnatural after being cropped and placed in the middle of a black background. ResMem might not be well-suited to discern the memorability differences of these unnatural animal images, as suggested by the smaller standard deviation of ResMem HR for animals (*SD* = .05), compared to scenes (*SD* = .11). This resonates with the ceiling effect in ResMem predictions for animals (min = .80, max = .95), suggesting ResMem’s incapability in capturing memorability differences between unnatural animal images.

One surprising result that requires further exploration was that experimentally-measured adults’ memory was not predictive of children’s memory. This is counterintuitive because ResMem could predict both adult and children’s memories, making us initially expect adults’ memory to also correlate with children’s memory. However, adults may have used more elaborate strategies or relied on extra visual information that ResMem does not capture, making adults’ memory a noisier model than ResMem in predicting children’s memory. With this, we can view ResMem as a useful model of children’s memorability and make predictions about children’s memories using the ResMem DNN without having to collect additional adult data. To further explore the lack of correlation between children’s and adults’ memories, a future step will be to measure children’s and adults’ memories using the same associative memory task or continuous recognition task. This is the first step to tease apart whether the lack of correlation between children’s and adults’ memories was due to task or strategy differences between the age groups.

### Concluding Remarks

This paper reveals the developmental trajectory of when children start to show consistencies in their visual memory. We provide evidence that children develop adult-like susceptibility to image memorability by the age of 4 through experience. Such susceptibility was the most salient when tested after a long, 1-week delay. Results suggest that the memorability effect emerges in early childhood through neural development or cognitive experience accumulation. These findings pave the way for exciting applications that could engage young children in more intuitive and memorable visual learning experiences. Teachers and caregivers could leverage the power of the ResMem DNN to select more memorable visual learning tools for children. Because young children at different ages seemed to use different visual memory strategies, education materials targeting young children could cater to different preferred visual strategies at different ages. We have outlined a few future directions of behavioral research to further investigate the development of visual memory. Remarkably, despite the individual differences that occur within and across age groups, by the age of 4, children already remember and forget the same images as adults.

