## Supplemental Materials for "Children Develop Adult-Like Visual Sensitivity to Image Memorability by the Age of Four"

301 Green Hall

5848 S. University Ave.

Chicago, IL 60637

Contents Page

**Supplemental Experiment S1:** Potential encoding strategies used by 3-year-olds. 3

**Supplemental Results:** Testing statistical power of the child within-age consistency analysis. 6

**Supplemental Results:** Splitting 4-year-olds using accuracy. 6

**Supplemental Results:** Results for logistic regression models that included interactions or adult HR for animals 7

**Supplemental Results:** Did children remember individual images or animal-scene associations? 8

**References** 11

**Supplementary Table 1.1:** $\beta$ estimates of the four logistic regression models 12

**Supplementary Table 1.2:** $\beta$ estimates of the four logistic regression models (using continuous ages) 14

**Supplementary Table 2:** Statistics of adult HR vs. children's memory accuracy 16

**Supplemental Experiment S1: Potential encoding strategies used by 3-year-olds.**

We found that ResMem was not able to predict the memory of 3-year-olds, but 3-year-olds were surprisingly consistent with one another in what images they remembered and forgot. We conjectured that 3-year-olds might have consistently used encoding strategies that were not adult-like and could not be captured by ResMem. One possibility is that 3-year-olds's familiarity with the type of place depicted in the scene image may affect their memories. Previous research noted that infants show selective attention to familiar shapes (Rose et al., 1982), faces, and objects (Roder et al., 2000) before attending to novel stimuli, and children remember familiar information better (Hudson & Nelson, 1986). The reason ResMem is not able to predict 3-year-olds' image memory may be that ResMem does not have access to information that is specifically familiar to 3-year-olds. For example, children at the age of 3 may be familiar with what playgrounds look like but not work offices, whereas adults have gained a broader range of visual experience including playgrounds and offices. With this, adults' memorability predicted by ResMem may not be able to capture memory biases based on what 3-year-olds are specifically familiar with. Another possibility for why ResMem did not predict 3-year-olds' memory may be that 3-year-olds better encode aesthetic images, whereas aesthetics and memorability have been found to be uncorrelated (Isola et al., 2014). Previous research found that infants look longer at more attractive adult (Samuels & Ewy, 1985) and infant faces (Van Duuren et al., 2003) as rated by adults, suggesting that adults' aesthetic ratings of images may capture the visual appeal of these images to infants. If 3-year-olds remembered scene images based on image aesthetics, ResMem-predicted memorability would fail to capture 3-year-olds' memory patterns. To find out whether the memory of infants at age 3 could be influenced by image familiarity and aesthetics, we asked adults to rate the 20 scene images from the child dataset on familiarity and visual appeal to 3-year-olds, and correlated adults' ratings with children's memory.

### Method

#### *Participants*

Thirty-one online participants (*N* Female = 17, *N* Male = 12, *N* non-binary/third gender = 2) on Prolific aged between 18 and 69 engaged in the rating task. To ensure the quality of participants, we recruited workers with approval rates of at least 90% and numbers of previous submissions of at least 50. We prescreened workers to have an IP address within the United States and to be fluent in English. We also excluded participants who answered any of the three common-sense questions incorrectly. We recruited 34 online participants to account for exclusion, and 31 adult participants were included for analysis upon applying the exclusion criteria.

***Materials and Procedure***

At the beginning of the experiment, participants agreed to the consent form approved by the Institutional Review Board (IRB19-1395) at the University of Chicago and answered three common-sense questions (e.g., "What is the name of the season after winter?") as an attention check. The main task consisted of the 20 scene images from the child dataset, and each was paired with two rating questions. The first question asked participants to rate the statement "this place is familiar to 3-year-olds" on a 5-point Likert scale from "highly disagree" to "highly agree." The second question followed the same format with the statement "this image is visually appealing to 3-year-olds." The study walked participants through these two rating questions in a practice trial before the main rating task to ensure participants' understanding. At the end of the experiment, participants received a demographic survey, which included basic demographic questions and three questions about their experience with 3-year-olds. Specifically, participants were asked to rate on a 5-point Likert scale (from "highly disagree" to "highly agree") for the statements "I have lots of experience living and working with 3-year-olds" and "I know a lot about 3-year-olds," and participants were asked whether they had young children. The full experiment took around 3 minutes. Participants received $0.40 upon completion, and they were allowed to end the experiment at any time.

**Results and Interpretations**

Before assessing whether 3-year-olds used image familiarity or aesthetics as encoding strategies, we ran a consistency analysis on each adult rating in this Supplemental Experiment to see whether adults consistently rated certain images as more familiar and attractive to 3-year-olds. We split adults into random halves and Spearman correlated the averaged familiarity or aesthetic ratings from each half over 100 iterations. We found that adults were highly consistent in their ratings of what images were more familiar ($M$ = 3.08, $SD$ = 1.02, averaged Spearman-

Brown corrected *rho* = .96, $p < .01$) and aesthetic ($M = 3.18$, $SD = .94$, averaged Spearman-Brown corrected *rho* = .96, $p < .01$) to 3-year-olds.

To see whether 3-year-olds used image familiarity or aesthetics as encoding strategies, we Spearman correlated the memory performance of 3-year-olds with familiarity or aesthetic ratings using the 20 scene images and found that neither predicted the memory of 3-year-olds (all $p > .05$). We then ran the same correlations between 3-year-olds' memory and ratings from adults who had more experience with infants. We found that the familiarity ratings from adults who had young children ($N = 6$, $M = 3.16$, $SD = .99$) significantly predicted the memory of 3-year-olds (*rho* = .59, $p = .01$), but the aesthetic ratings ($M = 3.08$, $SD = .76$) from the same adults did not ($p > .05$). Such results suggest that adults who are more experienced with young children might have more accurate judgments on children's visual experience and 3-year-olds likely prioritize familiar images during encoding. Another piece of supportive evidence was that the familiarity ratings from adults who had lots of experience living or working with 3-year-olds (rated "agree" or "highly agree") ($N = 9$, $M = 3.04$, $SD = 1.03$) significantly predicted the memory of 3-year-olds (*rho* = .49, $p = .03$), but the aesthetic ratings ($M = 3.28$, $SD = 1.02$) from the same adults did not ($p > .05$). Neither rating from adults who knew a lot about 3-year-olds (rated "agree" or "highly agree") ($N = 10$, familiarity: $M = 3.12$, $SD = .93$; aesthetics: $M = 3.18$, $SD = .85$) predicted 3-year-olds' memory (all $p > .05$). This suggests that having direct experience living or working with infants is crucial for making an accurate judgment on what places 3-year-olds are familiar with. These results suggest that the consistency in 3-year-olds' memory may have manifested in their tendency to utilize image familiarity but not aesthetics as a strategy to guide encoding.

We have found that 3-year-olds' experience of familiar places affects their memory of scene images. Further research could investigate whether there are other intrinsic properties of images, separate from the intrinsic memorability of those images, that affect the memory of young children, such as complexity.

### Supplemental Results

#### Testing statistical power of the child within-age consistency analysis.

We were curious to see whether the child within-age consistency analyses were potentially underpowered. The fact that 4- and 5-year-olds did not show consistent within-age memory patterns might be due to the low number of child participants upon splitting each age group into halves (i.e., approximately 22 participants per half) when running the split-halves consistency analyses. To see if the split led to low power, we split 4- and 5-year-olds into random halves, and Spearman correlated each with ResMem HR on the 20 scene images from the child dataset over 500 iterations. If the split were not underpowered, the average of the 1000 correlations within each age group would be significant because we previously found that ResMem could predict the memory of 4- and 5-year-olds when not downsampled to half the data. We found that, however, ResMem did not predict memory of either age group when split into halves (all $p > .05$, permutation tests). Indeed, 4- and 5-year-olds' inconsistent memory might be due to low power. Nevertheless, 3-year-olds showed robust consistency in memory even with low power.

#### Splitting 4-year-olds using accuracy.

An alternative way to probe the additional explanation that children might have developed adult-like encoding strategies around the age of 4 was to test whether 4-year-olds with better memories could be better predicted by ResMem than those with worse memories. Children's memory data showed that overall memory accuracy increased with age. We thus expected the high-accuracy group of 4-year-olds to show more adult-like memory patterns, i.e., remembering a certain set of images more than others. We performed a median split of 4-year-olds into a group with higher accuracy ($N = 22$, $M = .93$, $SD = .08$) and another with lower accuracy ($N = 22$, $M = .38$, $SD = .17$) across the eight test trials, and Spearman correlated each group with ResMem HR on the 20 scene images from the child dataset. Two scenes were excluded because they were never the target scenes for any test trials presented to the high-accuracy group of 4-year-olds, resulting in 18 scene images for this correlation analysis. We found that, however, ResMem could not predict the memory of 4-year-olds from either the high- or low-accuracy group (all $p > .05$). It is possible that grouping 4-year-olds based on birth months vs. memory accuracy may reveal separate age-related phenomena, such that older 4-year-olds showed more adult-like memory patterns but the high-accuracy group of 4-year-olds did not. In other words, it may be the age-related development in memory, rather than the improvement in memory performance, that causes children to take on adult-like patterns of memorability. The nonsignificant result in this section may also due to low power, as calculated in the previous section.

**Results for logistic regression models that included interactions or adult HR for animals**

Because we found that image memorability affected children's memory differently for different age groups and delay conditions, we tested a second logistic regression model that added two-way interaction terms of ResMem HR with age or delay to the base model:

$$children's\ memory\ accuracy \sim \beta_1(ResMem\ HR_{scenes}) + \beta_2(Age\ 4) + \beta_3(Age\ 5) + \beta_4(Delay\ 2) + \beta_5(Delay\ 3) + (1 \mid subject\ ID) + (1 \mid associated\ animal\ ID) + \beta_6(\boldsymbol{ResMem\ HR_{scenes} : Age\ 4}) + \beta_7(\boldsymbol{ResMem\ HR_{scenes} : Age\ 5}) + \beta_8(\boldsymbol{ResMem\ HR_{scenes} : Delay\ 2}) + \beta_9(\boldsymbol{ResMem\ HR_{scenes} : Delay\ 3})$$

None of the interaction terms were significant (all $p > .05$), and adding the interaction terms also drove all existing fixed effect predictors to be nonsignificant (all $p > .05$), except for the delay predictor for the difference between the shortest and the longest delays ($\beta_{Delay\ 3} = -2.62$, $p = .002$). Given the unchanged adjusted $R^2$ and increased AIC in the interaction model (adjusted $R^2 = .39$, $p < .001$, AIC = 5079) compared to the base model (adjusted $R^2 = .39$, $p < .001$, AIC = 5069), these results suggest that the base model better fits the children's memory data compared to the interaction model.

We tested a third model incorporating the animal memorability scores into the base model, by replacing the random intercept of associated animal ID in the base model with Animal HR (as predicted by ResMem):

$$children's\ memory\ accuracy \sim \beta_1(ResMem\ HR_{scenes}) + \beta_2(Age\ 4) + \beta_3(Age\ 5) + \beta_4(Delay\ 2) + \beta_5(Delay\ 3) + (1 \mid subject\ ID) + \beta_6(\boldsymbol{ResMem\ HR_{animals}})$$

The crucial added predictor, *ResMem Animal HR*, was nonsignificant ($p > .05$), suggesting that it was the target scene but not the associated cue animal image that drove children's memory. Similar to the base model, all other predictors were significant (all $p < .05$), but the adjusted $R^2$ decreased, and the AIC increased in the animal HR model (adjusted $R^2 = .38$, $p < .001$, AIC = 5073), suggesting the base model is a better fit (see Supplementary Table 1.1 for beta estimates). In conclusion, in comparison to the interaction and the animal HR models, the base model was optimal in predicting children's memory using ResMem HR of scenes, age, and delay. We ran the same logistic regression models using continuous rather than discrete ages (see Supplementary Table 1.2 for results) and found that the base model including discrete ages still led to the best model fit.

**Did children remember individual images or animal-scene associations?**

The child memory data was collected using an association task. In each trial, child participants encoded an animal image, a scene image, and a paired image, where the animal was placed in the scene. We were curious whether children remembered individual scene images or animal-scene-associated images. We ran Spearman correlations between children's memory ($M = .64$, $SD = .08$) and ResMem HR of individual scene images ($M = .73$, $SD = .11$) and animal-scene-paired images ($M = .82$, $SD = .07$). Note that in the child memory experiment (Saragosa-Harris et al., 2021), the same scene could be paired with different animals in different trials (for different children). Given that children's item-based scene memory was obtained by averaging across trials that involved the same scene, we obtained ResMem HR predictions for all animal-scene-paired images and averaged those that were associated with the same scene. With this, we obtained ResMem HR measures for each of the 20 scenes as they were involved in the animal-scene associations. There were 20 scene data points for children's memory, ResMem HR of individual scene images, and ResMem HR of animal-scene-paired images, respectively. We found that overall, ResMem HR of individual scenes could not predict children's memory (*rho* =.31, $p$ =.18), but ResMem HR of animal-scene combinations could (*rho* = .46, $p = .04$). This means that across ages and delay, children might have remembered the animal-scene-paired images rather than the individual scene images. We then ran Spearman correlations between ResMem HR of the paired images and children's memory in each age and delay group. Similar to the results of ResMem HR of individual scene images, ResMem HR for animal-scene-paired images significantly predicted the memory of 4 (*rho* = .48, $p = .03$) and 5-year-olds (*rho* = .58, $p = .007$) (FDR-corrected, $q < 0.05$), but not that of 3-year-olds ($p > .05$). Unlike the results of

ResMem HR of individual scene images, which could predict children's memory across age groups after a 1-week delay, ResMem HR of the paired images could not predict children's memory in any delay conditions (all $p > .05$). These more fine-grained results suggest that ResMem HR for individual images might be more robust in predicting children's memory, compared to ResMem HR of the animal-scene paired images.

Because we found mixed results from the overall vs. the fine-grained correlations, we ran a logistic regression (adjusted $R^2 = .37$, $p < .001$, AIC = 5056) following the base model introduced in the main paper with a slight alteration on the predictor of ResMem HR:

$$children's\ memory\ accuracy \sim \beta_1(\boldsymbol{ResMem\ HR_{animal-scene\ pairs}}) + \beta_2(Age\ 4) + \beta_3(Age\ 5) + \beta_4(Delay\ 2) + \beta_5(Delay\ 3) + (1 \mid subject\ ID)$$

The random intercepts of the paired animal ID were deleted from the model because *ResMem HR for animal-scene paired images* already accounted for the variation from the nature of the association task. We found that both age predictors were significant ($\beta_{Age\ 4} = .70$, $p = .02$; $\beta_{Age\ 5} = 1.29$, $p < .001$). Both delay predictors were also significant ($\beta_{Delay\ 2} = -.93$, $p = .002$; $\beta_{Delay\ 3} = -1.93$, $p < .001$). This means that the differences between ages and delays explain variations in children's memory, as in the base model. This model had a lower adjusted $R^2$ but also a better (lower) AIC compared to the base model (adjusted $R^2 = .39$, $p < .001$, AIC = 5069). However, the crucial predictor, *ResMem HR of the animal-scene paired images*, was nonsignificant ($\beta_{ResMem\ HR\ animal\text{-}scene\ pairs} = 1.68$, $p = .07$), suggesting that the animal-scene paired images did not drive children's memory. We also ran a model that replaced the discrete age predictors with a continuous age predictor (see Supplementary Table 1.2 for results) and found a worse model fit and $\beta$ estimates closer to zero. Hence, we have mainly discussed the model with discrete age predictors here. Despite the similarity between scene images and animal-scene-paired images, illustrated by a significant Spearman correlation between children's memory of the scene vs. animal-scene-paired images (*rho* = .69, $p = .001$), the target scene seems to be the main driving force of children's memory.

We found that taking the memorability of the associated animal into consideration did not greatly improve predictions of children's memory. More evidence is needed to disambiguate children's encoding processes. A future direction is to directly test the memorability of image associations among children and compare the results to children’s memorability patterns of individual images.

**Supplementary Table 1.1**

*$\beta$ estimates of the four logistic regression models*

| $\beta$ Estimates | Base Model | Interaction Model | Animal HR Model | Animal-Scene Association Model |
|---|---|---|---|---|
| $ResMem\ HR_{scenes}$ | .85* | -.02 | .90** | — |
| $ResMem\ HR_{animals}$ | — | — | -.28 | — |
| $ResMem\ HR_{animal-scene\ pairs}$ | — | — | — | 1.68 |
| $Age$ 4 | .70* | -.20 | .70* | .70* |
| $Age$ 5 | 1.31*** | .11 | 1.31*** | 1.29*** |
| $Delay$ 2 | -.94** | -.58 | -.94** | -.93** |
| $Delay$ 3 | -1.95*** | -2.62** | -1.95*** | -1.93*** |
| $ResMem\ HR_{scenes}$ : $Age$ 4 | — | .97 | — | — |
| $ResMem\ HR_{scenes}$ : $Age$ 5 | — | 1.30 | — | — |
| $ResMem\ HR_{scenes}$ : $Delay$ 2 | — | -.38 | — | — |
| $ResMem\ HR_{scenes}$ : $Delay$ 3 | — | .73 | — | — |
| $adjusted\ R^2$ | .39 | .39 | .38 | .37 |
| $p$ | <.001 | <.001 | <.001 | <.001 |
| AIC | 5069 | 5079 | 5073 | 5056 |

*Note*. Formulas for these four logistic regression models (from the main paper and the supplemental materials) are shown below.

*$p$ < .05. **$p$ < .01. ***$p$ < .001.

**Base Model:**

$$children's\ memory\ accuracy \sim \beta_1(ResMem\ HR_{scenes}) + \beta_2(Age\ 4) + \beta_3(Age\ 5) + \beta_4(Delay\ 2) + \beta_5(Delay\ 3) + (1 \mid subject\ ID) + (1 \mid associated\ animal\ ID)$$

**Interaction Model:**

$$children's\ memory\ accuracy \sim \beta_1(ResMem\ HR_{scenes}) + \beta_2(Age\ 4) + \beta_3(Age\ 5) + \beta_4(Delay\ 2) + \beta_5(Delay\ 3) + (1 \mid subject\ ID) + (1 \mid associated\ animal\ ID) + \beta_6(\boldsymbol{ResMem\ HR_{scenes} : Age\ 4}) + \beta_7(\boldsymbol{ResMem\ HR_{scenes} : Age\ 5}) + \beta_8(\boldsymbol{ResMem\ HR_{scenes} : Delay\ 2}) + \beta_9(\boldsymbol{ResMem\ HR_{scenes} : Delay\ 3})$$

**Animal HR Model:**

$$children's\ memory\ accuracy \sim \beta_1(ResMem\ HR_{scenes}) + \beta_2(Age\ 4) + \beta_3(Age\ 5) + \beta_4(Delay\ 2) + \beta_5(Delay\ 3) + (1 \mid subject\ ID) + \beta_6(\boldsymbol{ResMem\ HR_{animals}})$$

**Animal-Scene Association Model:**

$$children's\ memory\ accuracy \sim \beta_1\left(\boldsymbol{ResMem\ HR_{animal-scene\ pairs}}\right) + \beta_2(Age\ 4) + \beta_3(Age\ 5) + \beta_4(Delay\ 2) + \beta_5(Delay\ 3) + (1 \mid subject\ ID)$$

**Supplementary Table 1.2**

*$\beta$ estimates of the four logistic regression models (using continuous age)*

| $\beta$ Estimates | Base Model | Interaction Model | Animal HR Model | Animal-Scene Association Model |
|---|---|---|---|---|
| $ResMem\ HR_{scenes}$ | .84* | -.1.43 | .89** | — |
| $ResMem\ HR_{animals}$ | — | — | -.28 | — |
| $ResMem\ HR_{animal-scene\ pairs}$ | — | — | — | .63* |
| $Continuous\ Age$ | .68*** | .24 | .67*** | .67*** |
| $Delay\ 2$ | -.86** | -.54 | -.85** | -.85** |
| $Delay\ 3$ | -1.92*** | -2.59** | -1.92*** | -1.90*** |
| $ResMem\ HR_{scenes}$ $:Continuous\ Age$ | — | .48 | — | — |
| $ResMem\ HR_{scenes} : Delay\ 2$ | — | -.34 | — | — |
| $ResMem\ HR_{scenes} : Delay\ 3$ | — | .72 | — | — |
| $adjusted\ R^2$ | .39 | .39 | .39 | .37 |
| $p$ | <.001 | <.001 | <.001 | <.001 |
| AIC | 5071 | 5081 | 5074 | 5061 |

*Note*. Formulas for these four logistic regression models (replace discrete *Age* predictors in models from Supplementary Table 1.1 using *Continuous Age*) are shown below.

*$p$ < .05. **$p$ < .01. ***$p$ < .001.

**Base Model:**

$$children's\ memory\ accuracy \sim \beta_1(ResMem\ HR_{scenes}) + \beta_2(\boldsymbol{Continuous\ Age}) + \beta_3(Delay\ 2) + \beta_4(Delay\ 3) + (1 \mid subject\ ID) + (1 \mid associated\ animal\ ID)$$

**Interaction Model:**

$$children's\ memory\ accuracy \sim \beta_1(ResMem\ HR_{scenes}) + \beta_2(\boldsymbol{Continuous\ Age}) + \beta_3(Delay\ 2) + \beta_4(Delay\ 3) + (1 \mid subject\ ID) + (1 \mid associated\ animal\ ID) + \beta_5(\boldsymbol{ResMem\ HR_{scenes} : Age\ 4}) + \beta_6(\boldsymbol{ResMem\ HR_{scenes} : Age\ 5}) + \beta_7(\boldsymbol{ResMem\ HR_{scenes} : Delay\ 2}) + \beta_8(\boldsymbol{ResMem\ HR_{scenes} : Delay\ 3})$$

**Animal HR Model:**

$$children's\ memory\ accuracy \sim \beta_1(ResMem\ HR_{scenes}) + \beta_2(\boldsymbol{Continuous\ Age}) + \beta_3(Delay\ 2) + \beta_4(Delay\ 3) + (1 \mid subject\ ID) + \beta_5(\boldsymbol{ResMem\ HR_{animals}})$$

**Animal-Scene Association Model:**

$$children's\ memory\ accuracy \sim \beta_1(\boldsymbol{ResMem\ HR_{animal-scene\ pairs}}) + \beta_2(\boldsymbol{Continuous\ Age}) + \beta_3(Delay\ 2) + \beta_4(Delay\ 3) + (1 \mid subject\ ID)$$

**Supplementary Table 2**

*Statistics of adult HR vs. children's memory accuracy across ages or delays*

| Children's Memory Accuracy | | | Adult HR | |
|---|---|---|---|---|
| *M* | *SD* | | *M* = .73 | *SD* = .11 |
| | | | *rho* | *p* |
| .52 | .19 | 3 years old | .10 | .66 |
| .62 | .15 | 4 years old | .05 | .83 |
| .76 | .07 | 5 years old | .13 | .58 |
| .79 | .09 | 5-min delay | -.09 | .71 |
| .66 | .13 | 24-h delay | .17 | .48 |
| .47 | .11 | 1-week delay | .41 | .07 |

*Note*. Statistics for the Spearman correlations between Adult HR for scenes and children's memory accuracy at each age across delays and at each delay across ages.
